# Calcium entry does not drive fast mechanotransduction adaptation in cochlear hair cells

**DOI:** 10.1101/629626

**Authors:** Giusy A. Caprara, Andrew A. Mecca, Yanli Wang, Anthony J. Ricci, Anthony W. Peng

## Abstract

Sound detection in auditory sensory hair cells depends on the deflection of the stereocilia hair bundle, which opens mechano-electric transduction (MET) channels. Adaptation is hypothesized to be a critical property of MET that contributes to the wide dynamic range and sharp frequency selectivity of the auditory system. Historically, adaptation was hypothesized to have multiple mechanisms, all of which require calcium entry through MET channels. Our recent work using a stiff probe to displace hair bundles showed that the fastest adaptation mechanism (fast adaptation) does not require calcium entry. Using a fluid-jet stimulus, others obtained data showing only a calcium-dependent fast adaptation response. Here, we identified the source of this discrepancy. Because the hair cell response to a hair bundle stimulus depends critically on the magnitude and time course of the hair bundle deflection, we developed a high-speed imaging technique to quantify this deflection. The fluid jet delivers a force stimulus, and step-like force stimuli lead to a complex time course of hair bundle displacement (mechanical creep), which affects the hair cell’s macroscopic MET current response by masking the time course of the fast adaptation response. Modifying the fluid-jet stimulus to generate a step-like hair bundle displacement produced rapidly adapting currents that did not depend on membrane potential. This indicated that fast adaptation does not depend on calcium entry. We also confirmed the presence of a calcium-dependent slow adaptation process. These results confirm the existence of multiple adaptation processes: a fast adaptation that is not driven by calcium entry and a slower calcium-dependent process.

**Significance Statement:** Mechanotransduction by sensory hair cells represents a key first step for the sound sensing ability in vertebrates. The sharp frequency tuning and wide dynamic range of sound sensation are hypothesized to require a mechanotransduction adaptation mechanism. For decades, it had been accepted that all adaptation mechanisms require calcium entry into hair cells. However, more recent work indicated that the apparent calcium dependence of the fastest adaptation differs with the method of cochlear hair cell stimulation. Here, we reconcile existing data and show that calcium entry does not drive the fastest adaptation process, independent of the stimulation method.

## Introduction

Inner ear hair bundles comprise an array of graded length stereocilia that are organized in a staircase manner. Positive hair bundle deflection increases tension in a filamentous tip link that connects adjacent rows of stereocilia (1–3). Changes in tip-link tension result in the gating of mechano-electric transduction (MET) ion channels.

MET adaptation is hypothesized to extend the dynamic range of the cell, control the channel’s operating point, and filter incoming stimuli (4–6). Adaptation is identified by two phenomena: 1) a decay in current amplitude (“current decay”) during a sustained hair bundle displacement (5–8), and 2) a shift in the operating point (i.e., position along the displacement axis) of the MET channel’s current vs. displacement curve (“activation curve”), without a change in the channel’s sensitivity (i.e., the activation curve slope) during a constant displacement step, hereafter called adaptation shift (5–7, 9). The former is assayed using displacement steps and the latter is assayed using two-pulse experiments, where an activation curve is generated before and after a sustained displacement. In non-mammalian vertebrates, at least two adaptation processes have been described, with different decay time constants, different extents, and different calcium sensitivities (10–13). Slow adaptation has time constants of 10-100 ms and is hypothesized to be a calcium-dependent modulation of tip-link tension via myosin motor molecules controlling the position of the upper tip-link insertion site (14, 15). Fast adaptation has a time constant of ~1 ms or less. Multiple potential mechanisms are suggested for fast adaptation, all of which involve calcium entering through MET channels and binding to an intracellular site that is either the channel itself or very close to it (6, 14, 16-20).

Several experimental results from non-mammalian vertebrates support a model in which calcium must enter the hair cell to affect both fast and slow adaptation: 1) Current decay occurs at negative potentials where calcium entry occurs, but not at positive potentials where calcium entry is inhibited (6, 21, 22). 2) Two-pulse experiments demonstrate adaptation shifts at negative potentials that are modulated when using different extracellular calcium concentrations (5–7). 3) The time constants of fast and slow adaptation increase with increasing intracellular calcium buffering or decreasing extracellular calcium concentrations (5, 6, 11, 20, 23). 4) The resting open probability of MET channels, an indicator of the activation curve operating point, increases with lower extracellular calcium concentration and upon depolarization (6, 11, 20, 24). These data have been interpreted as supporting calcium-dependent adaptation mechanisms. Our recent data from cochlear hair cells challenge this hypothesis for mammalian auditory hair cells (12, 13, 25).

For *in vitro* MET experiments, stiff probes and fluid jets are the most commonly used methods; they deliver step-like displacement or force stimuli to the hair bundle, respectively. Stiff probes directly couple to stereocilia. In contrast, fluid jets eject extracellular solution onto hair bundles from the tip of a pipette; the fluid velocity produces a drag force stimulus on the hair bundle (26).

Our previous experiments with mammalian cochlear hair cells used stiff probe stimuli (12, 13). Improvements to stiff probe technology produced step-like displacement rise times as short as 11 µs (12), which provided finer temporal resolution of channel activation and adaptation properties. With this technology, we found that fast adaptation does not slow with depolarization or lower extracellular calcium and that adaptation shifts are not affected by depolarization, suggesting that fast adaptation is not driven by calcium entry (12). We performed multiple control experiments to rule out stimulus artifacts (27) that might have been interpreted as fast adaptation (12). Furthermore, we found that changes in the resting MET channel open probability induced by lowering extracellular calcium or depolarization (experimental evidence 4) result from a lipid modulation mechanism that is separate from fast adaptation and occurs with a time constant of ~125 ms (13). Therefore, changes in MET channel resting open probabilty alone cannot be used to support the calcium dependence of fast adaptation. These data supported that fast adaptation was not driven by calcium entry.

In contrast, data acquired by other labs using fluid-jet stimulators with longer rise times of ~500 µs suggest that fast adaptation is driven by calcium entry (28). The goal of the present work is to reconcile these disparate findings.

We used a novel high-speed imaging system to quantify the magnitude and time course of fluid-jet stimuli and the resulting hair bundle displacement. With step-like force stimuli, the fast phase of the current decay was not observed at any potential. However, with fluid-jet stimuli modified to produce step-like hair bundle displacements, similar to what a stiff probe generates, we observed fast current decays; however, this did not depend on the membrane potential. These results are consistent with our stiff probe data and demonstrate that calcium entry does not drive the fast adaptation response.

## Results

### Mechanical creep in hair bundle displacement

Stiff probe and fluid-jet stimulations are inherently different; the former delivers a displacement and the latter a force. With the fluid jet, the resulting stereocilia displacement elicited will vary depending on the mechanical properties of the stereocilia. Thus, if the hair bundle dynamically changes its compliance, the resulting hair bundle displacement will also change dynamically. Previous experiments measured hair bundle movement using a photodiode system (14, 29, 30) or calibrated the fluid-jet driver voltage to a hair bundle displacement (28, 31, 32). The photodiode method can only image a portion of a hair bundle and assumes that the hair bundle moves as a unit. Additionally, the photodiode method is experimentally difficult for two major reasons: 1) the hair bundle needs to be precisely aligned with the photodiode, and 2) each experiment requires a calibration to convert photodiode voltage to displacement. Therefore, displacement measurements are often not done for all experiments. The fluid-jet driver voltage calibration method assumes that hair bundle mechanical properties (i.e., compliance) do not change for each stimulus paradigm or across different cells and that the same force always results in the same hair bundle displacement. However, mechanical properties are well-documented to change with voltage and during stimulation (14, 33, 34), therefore invalidating this assumption.

To measure hair bundle displacement in every experiment, we developed a novel method to quantify cochlear hair bundle displacement using high-speed imaging with image analysis. Imaging at speeds up to 50,000 frames per second allowed tracking of the motion of the whole hair bundle in all directions in the plane of imaging. This technique is experimentally easier because: 1) only a one-time calibration of the size of each pixel is required, and 2) no precise hair bundle alignment is required. To validate our imaging technique, we compared the high-speed imaging technique with the photodiode technique using a flexible glass fiber attached to a piezo-electric actuator. We found similar results between the two methodologies (Fig. 1A). By allowing the stimulated glass fiber to resonate, we demonstrated that the imaging approach had both the speed and sensitivity necessary to track nanometer scale displacement on a sub-millisecond timescale (Fig. 1A, bottom).

**Fig. 1.**
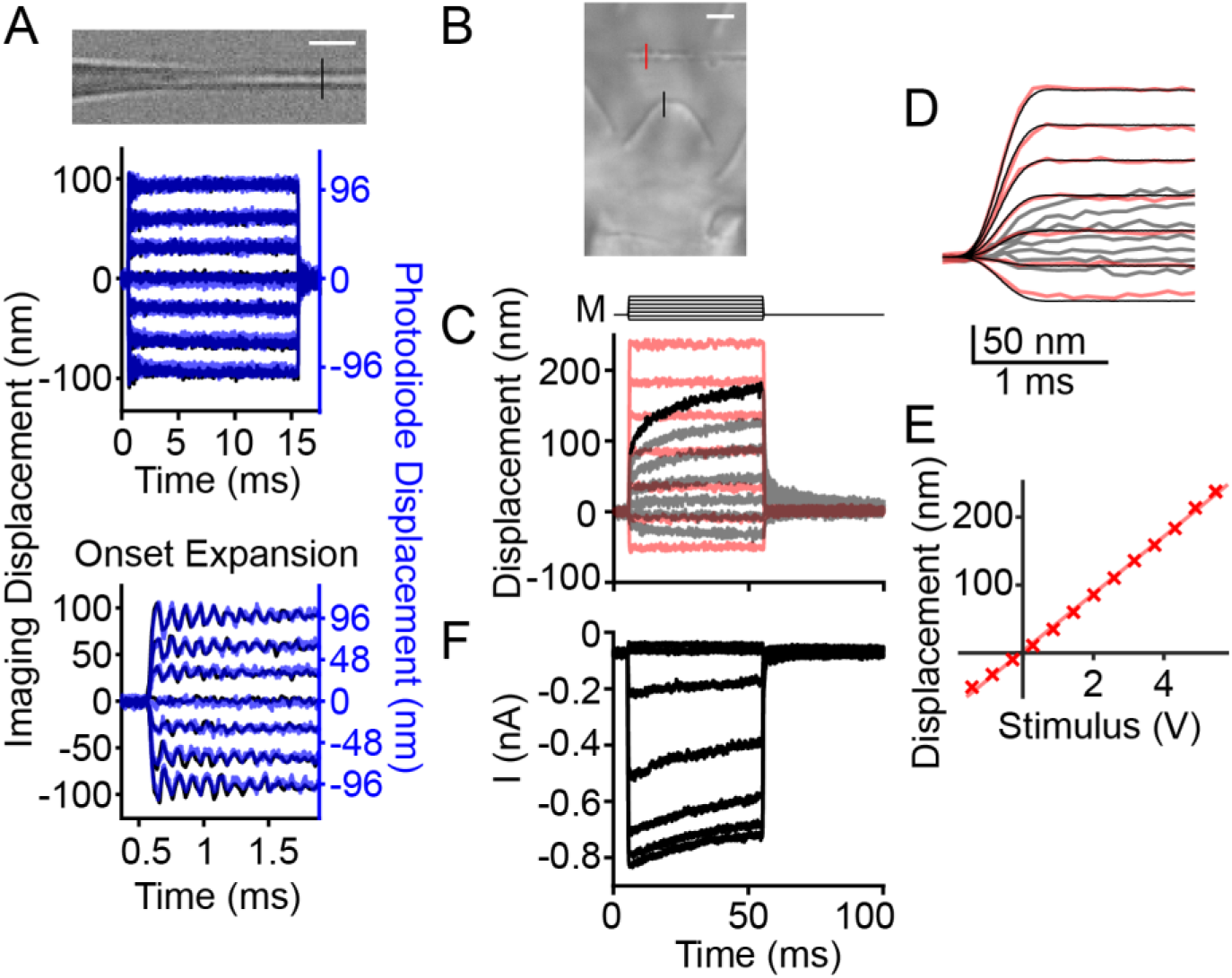
Validation of imaging and stimulus techniques. (A) High-speed imaging of an actuated flexible fiber at 50,000 frames per second (black traces) and the classical photodiode technique (blue, semi-transparent) yielded similar results in fiber displacement. An expansion of the onset of the fiber displacement showed an oscillation of ~9 kHz with both techniques. (B) Image of a hair bundle is shown with a passive flexible fiber in view to serve as a readout of the force applied by the fluid jet. Colored lines indicate where motion was measured. (C) In a single stimulation protocol, we monitored the fluid-jet force with the flexible fiber displacement (red), and we measured hair bundle displacement (gray). The hair bundle has a continued movement in the direction of the stimulation (i.e., mechanical creep, highlighted in one trace in black) even when the force from the fluid jet was step-like. M indicates the fluid-jet stimulus voltage waveform. (D) A time expansion of the onset of the fiber and hair bundle displacements from C shows the short rise time of the force step. The stimulus waveform (M, delayed and normalized) is overlaid in black to show that the stimulus voltage waveform kinetics match the force output. (E) The fiber displacement plotted against the stimulus voltage indicates linearity within the stimulus range used. (F) From the experiment in panel C, the resulting MET current at −84 mV holding potential shows the presence of slow current decay and the absence of fast current decay. Scale bars = 2 µm.

To confirm that the fluid-jet stimulator delivered a fast and step-like force, we used the tip displacement of a non-actuated flexible fiber as a readout of the fluid-jet force kinetics (Fig. 1B). The fiber moved as a step rising to a plateau in 0.5 ms, indicating that the fluid-jet force (Fig. 1C,D, red) had the same onset kinetics as the stimulus voltage (Fig. 1C, M traces; Fig. 1D, black traces). We also confirmed that force varied linearly with driver voltage by plotting the fiber displacement against the stimulus voltage (Fig. 1E). Despite a step-like fiber displacement, the hair bundle in the same field of view exhibited a mechanical creep (Fig. 1C, highlighted black trace), which is a continued movement in the direction of the applied force. This creep indicates an increasing hair bundle compliance as observed by others (14, 34-36). For the largest stimulation in 46 recorded cells, the creep was fit with a double exponential decay with time constants of τ_1_ = 1.8 ± 0.8 ms and τ_2_ = 20.4 ± 9.1 ms with the slower time constant contributing 56.2 ± 6.9% of the total creep. The recorded MET current elicited by fluid-jet force steps did not show fast current decays (defined as exponential decay time constant < 5 ms) but did show slow current decays (Fig. 1F). For step sizes eliciting ~50% of the maximum current in the same 46 cells, current decays fit with a single exponential decay had τ = 26.5 ± 21.0 ms. These data indicate that step-like force stimuli exhibit hair bundle creep and lack fast current decay.

We excluded the possibility that the observed creep was an artifact of underlying epithelium movement by performing the same displacement analysis on features of the epithelial surface below the hair bundle (Fig. S1). Changes in displacement from 0.5 ms after stimulus onset to the end of the force step for the epithelial surface were much smaller than the hair bundle creep for the same stimulus intensity (Fig. S1B). These data suggest the creep was not an artifact of epithelial movement.

### Fast current decay present with overshoot stimuli only at negative potentials

Unlike MET currents previously elicited with fluid-jet stimuli (28, 32), our MET currents did not show a fast current decay (Fig. 1F). Upon detailed inspection of the data from Corns et al. (2014), we found that they applied an underdamped step response, which is characterized by an overshoot and ringing of their step stimulus. Both overshoot and ringing are visible in their data with the carbon fiber. They used two carbon fibers to characterize their force stimulus. The first fiber (8 µm diameter, see Corns et al. (2014) Supplemental Fig. S2A) shows a step-like response with little to no evidence of an underdamped response. However, this fiber has a rise time of ~2.5 ms, much longer than the expected rise time of ~0.5 ms based on their stimulus voltage waveform. This indicates that viscous drag on their long (>2 mm) carbon fibers filtered the force readout. They also used a 10 µm diameter fiber with a shorter rise time of ~1.9 ms, indicating less filtering and the ability to observe more of the stimulus kinetics. This fiber displacement clearly shows an underdamped response (see Corns et al. (2014) Supplemental Fig. S2B). If they had a faster readout of their force stimulus with a shorter rise time of 0.5 ms (i.e., even less filtering of the response), they would likely observe an even larger initial overshoot. The possibility that their stimulus driving circuit produced ringing and that the fluid movement itself had ringing is further supported by the small oscillations observed in their MET currents and hair bundle displacements whose kinetics match the oscillations observed in their fiber displacement (see Corns et al. (2014) Supplemental Fig. S3). We sought to test whether an underdamped stimulus could account for the fast current decays observed by Corns et al. at negative potentials.

Based on the hair bundle displacement time course observed with our fluid-jet stimuli, we predicted that an overshoot of a force step would result in an early plateau of the hair bundle displacement. The fluid-jet stimulus used by Corns et al. (2014) led to an early plateau in the hair bundle displacement (Fig. 2A arrow, black reproduced from Corns et al. (2014)). We mimicked this key feature of the underdamped response to achieve a similar early plateau in the hair bundle displacement (Fig. 2A, blue) by simply using an overshoot stimulus with a single exponential decay back to steady state (Fig. 2B, arrow). Comparing our hair bundle displacement resulting from a step-like force to that of Corns et al. (2014) also highlights the ringing in their hair bundle displacement (Fig. 2A, magenta vs. black, arrowheads). Using an overshoot stimulation, we were able to reproduce two key results observed by Corns et al. (2014) regarding fast adaptation. 1) Fast current decays, which look similar to current decays caused by fast adaptation, were observed at negative potentials for positive displacements and upon returning from a negative displacement (Fig. 2B, arrowheads). 2) Both of these fast current decays were absent at positive potentials (Fig. 2B). When returning to a step-like force stimulus in the same cell, we observed our typical responses where no fast current decays were present (Fig. 2C). Interestingly, these results show that a small difference in the hair bundle displacement (Fig 2A, blue vs. magenta) led to a large difference in the MET current (Fig. 2B,C), which reinforces the hair bundle’s sensitivity to small displacement changes. These results indicate that the fast current decays observed by Corns et al. (2014) could be an artifact of an underdamped stimulus and not a manifestation of fast adaptation.

**Fig. 2.**
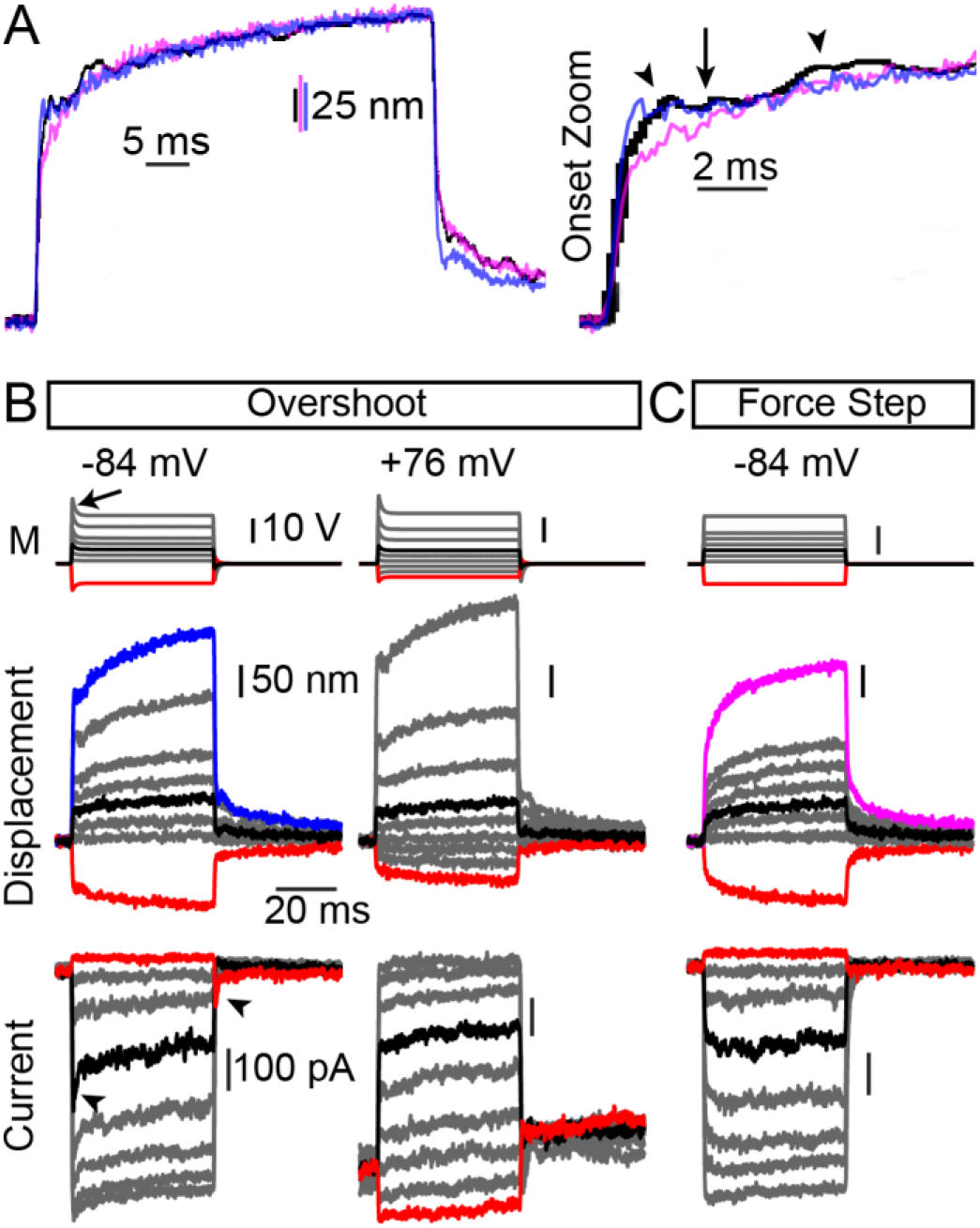
Overshoot stimuli can account for fast current decays observed by Corns et al. (2014). (A) To mimic the displacement responses of Corns et al. (2014) (black traces, reproduced from Corns et al. (2014)), we modified our step-like force stimulus (resulting displacement in magenta) to achieve an early plateau (arrow in onset zoom) in the displacement response using an overshoot stimulus (resulting displacement in blue). An expansion of the onset is shown to the right with the early plateau (arrow) and the oscillations (arrowheads) in the black trace indicated. (B) The stimulus waveform, resulting hair bundle displacements, and the recorded currents are plotted. Using 1 mM intracellular BAPTA buffer, overshoot stimuli resulted in current decays that appear like fast adaptation (arrowheads). In the same cell, the fast current decays were absent at positive potentials with overshoot stimuli. (C) When removing the overshoot and using a step-like force stimulation in the same cell, no fast current decays are evident at negative potentials. No averaging of stimulations was used, and current traces were 5 point smoothed using MATLAB’s smooth function; displacement traces were averaged over 14 lines around the apex of the hair bundle.

### Adaptation shifts indicate an adaptation mechanism that does not require calcium entry

Measuring adaptation shifts in response to a constant adapting step with a two-pulse protocol assesses adaptation (5, 12). Our experiments with a stiff probe in cochlear hair cells using two-pulse protocols found that the adaptation shifts for 5 ms sustained displacements were the same at negative and positive potentials (12), while others using a fluid jet found that adaptation shifts for 20 ms of sustained force stimuli were present at negative potentials but absent at positive potentials (28). The fluid-jet data did not have corresponding hair bundle displacement with each fluid-jet experimental paradigm; therefore, the data did not account for the mechanical creep (see Corns et al. (2014) Methods, last paragraph). Although the results are presented as current vs. displacement plots, they represent current vs. stimulus voltage plots. We performed these experiments while measuring hair bundle displacement using the high-speed imaging system, allowing us to simultaneously generate current vs. stimulus voltage and current vs. displacement plots. We used a three-pulse protocol where we assayed the adaptation shifts at 10 ms and 50 ms after the start of a sustained force step with 5 ms test pulses to generate the activation curve (Fig. 3A, M traces). We recapitulated the findings of Corns et al. (2014) with current vs. stimulus voltage plots (Fig. 3A, B), where a shift occurred at negative but not positive potentials. Additionally, current decay was observed at negative, but abolished at positive potentials (Fig. 3A, arrowheads). The current decay at negative potentials during the first 10 ms of the sustained force stimulus was fit with a single exponential decay τ = 14.0 ± 1.5 ms (n = 6), a time scale consistent with slow adaptation. These results could be interpreted as only calcium-dependent adaptation mechanisms.

**Fig. 3.**
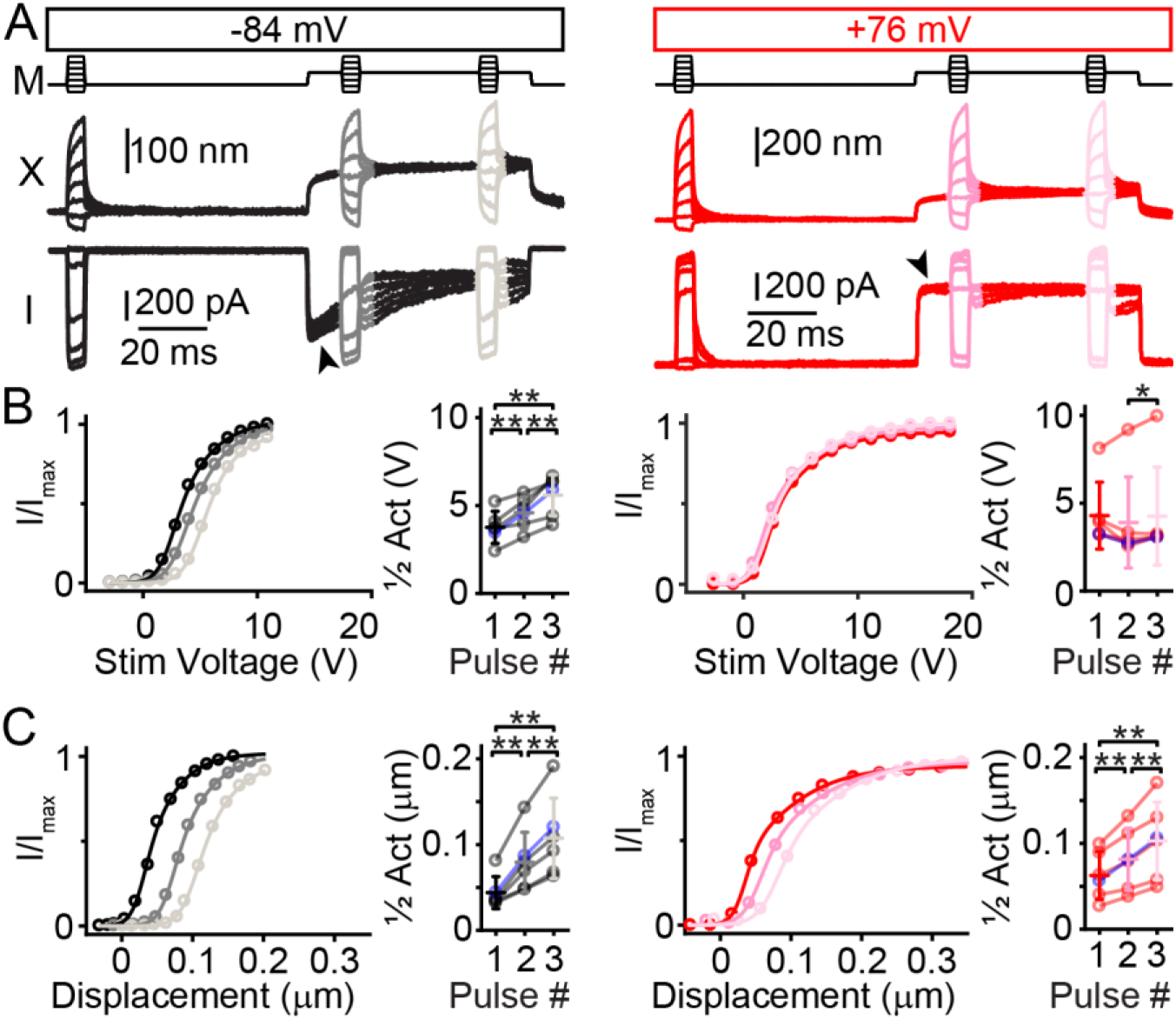
Three-pulse experiments to assay adaptation shifts using a fluid-jet stimulus and 0.1 mM BAPTA intracellular solution. (A) Voltage commands for the fluid jet in a three-pulse protocol are shown (M) at −84 mV (black) and at +76 mV (red) holding potentials with measured hair bundle displacements (X) and currents (I). Arrowheads indicate current decay at negative, but not positive potentials. (B) Analyzing data with the stimulus voltage showed activation curve shifts at negative potentials, but not positive potentials. Summary data of half activation points for each stimulus pulse show positive adaptation shifts at negative (n=6, pulse 1-2 *p*=0.0029, 2-3 *p*=0.0066, and 1-3 *p*=0.0039), but not positive potentials (n=6, pulse 1-2 *p*= *p*=0.28, 2-3 *p*=0.034, and 1-3 *p*=0.94). Blue data points are from traces in A. (C) Analyzing data with measured hair bundle displacement indicates adaptation shifts at both negative (n=6, pulse 1-2 *p*=0.0050, 2-3 *p*=0.0021, and 1-3 *p*=0.0033), and positive potentials (n=6, pulse 1-2 *p*=0.0045, 2-3 *p*=0.0042, and 1-3 *p*=0.0039).

In contrast, plotting the same currents against measured hair bundle displacement demonstrated adaptation shifts at both negative and positive potentials (Fig. 3C). These data suggested that adaptation shifts and current decays were decoupled when using sustained force stimuli because adaptation shifts occurred in the absence of current decays at positive potentials. Unlike our data with stiff probes (12), adaptation shifts with the fluid jet were significantly larger at negative potentials (35 ± 18 nm at 10 ms and 63 ± 30 nm at 50 ms, n = 6) than at positive potentials (19 ± 9 nm and 40 ± 19 nm at 50 ms, n = 6; *p*=0.035 at 10 ms and *p*=0.023 at 50 ms). These data could be interpreted in two ways with regard to the calcium dependence of adaptation. One interpretation is that since adaptation shifts occurred at positive potentials, there is an adaptation process that is not driven by calcium entry. That the adaptation shifts were larger at negative potentials as compared to positive potentials would indicate that there is a second calcium-dependent adaptation process; this could be the slow adaptation studied by others with a time course of ~10 ms (5, 21). An alternative interpretation is that all the adaptation observed at positive potentials is residual calcium-dependent slow adaptation. The motor model of slow adaptation does not preclude slow adaptation from occurring at positive potentials, since only the rate of adaptation is modulated by calcium entry (15). Although this is theoretically possible for the motor model, it has not been observed in non-mammalian hair cells, as no slow current decays resulting from displacement steps are observed during depolarization (6, 20, 21, 24). Here, the adaptation shift at positive potentials after 10 ms was 47.5% of the shift after 50 ms. If all of the adaptation shift was due to slow adaptation, then based on the single exponential decay equation, the time constant of the slow adaptation at positive potentials would need to be 16.7 ms (see methods for calculation), similar to the slow adaptation time constant at negative potentials of 14 ms. Since the time constants would be similar at negative and positive potentials, this would not fit with the motor model of slow adaptation, which predicts that the rate of adaptation at positive potentials is much slower than at negative potentials (15, 24). Therefore, the more likely interpretation is the existence of two adaptation mechanisms that differ in calcium dependence: a fast adaptation that does not require calcium entry and a slow adaptation that depends on calcium.

### Step-like displacement stimulations with the fluid jet unmask fast current decays

To further support the existence of two adaptation mechanisms with different calcium requirements, we tested if fast current decays could be observed at positive potentials using step-like displacement stimuli. We wondered if the fast current decays were masked by the hair bundle creep (Fig. 1C), where continued hair bundle movement counteracted the current decay and would be responsible for the lack of visible fast current decays with step-like force stimuli. To test this, we designed a circuit to modify the fluid-jet voltage waveform to create a step-like displacement of the hair bundle (Fig. 4A). To limit the contribution of calcium-dependent adaptation mechanisms, we used a high intracellular calcium buffer (10 mM BAPTA) in these experiments. We applied both step-like force and step-like displacement stimuli on the same cell for these experiments. Consistent with Figs. 1F and 2C, fast current decays were absent at both negative and positive potentials with step-like *force* stimuli (Fig. 4A). In the same cell with step-like *displacement* stimuli, fast current decays were present at both negative and positive potentials (Fig. 4A, arrowheads). For displacement steps eliciting ~50% maximum current, the fast current decay was similar at negative and positive potentials: at negative potentials it had a time constant of 1.6 ± 1.2 ms and contributed 34 ± 14% (n = 7) of the total extent of adaptation, and at positive potentials it had a time constant of 1.8 ± 1.4 ms and contributed 29 ± 17% (n = 6) of the total extent of adaptation (*p* = 0.99 and *p* = 0.32, respectively; paired Student’s t-test between negative and positive potentials). The slow current decay, on the other hand, was faster at negative potentials (14 ± 3 ms; n = 7) than at positive potentials (59 ± 41 ms; n = 6; *p* = 0.053, paired Student’s t-test). These data further support that all adaptation at positive potentials cannot be accounted for by residual slow adaptation, since a measured time constant of 1.8 ± 1.4 ms (n = 6) (Fig. 4A) is too fast for slow adaptation. The fast current decays at both negative and positive potentials (Fig. 4A, arrowheads) supported the conclusion that fast adaptation was not driven by calcium entry.

**Fig. 4.**
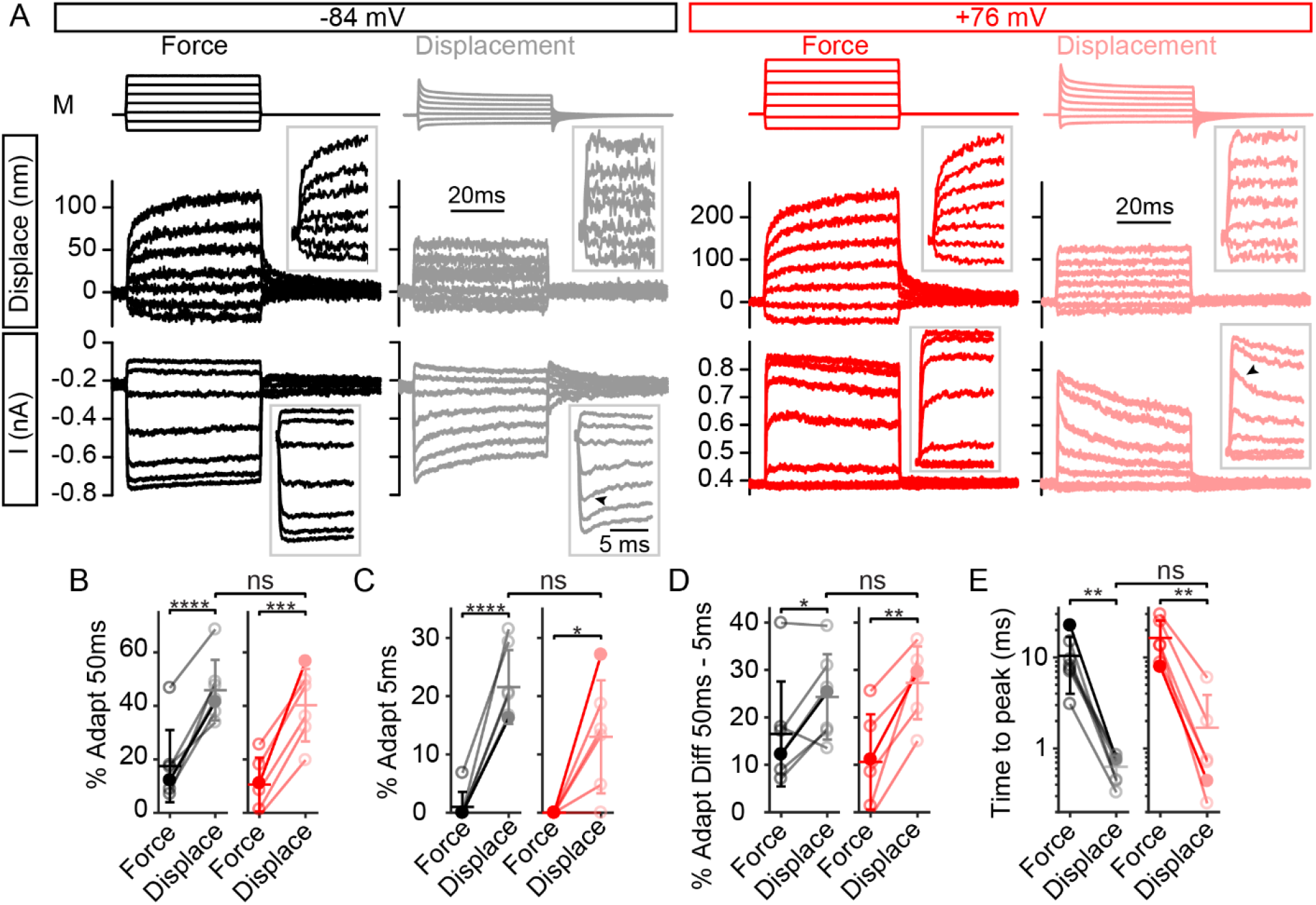
Step-like hair bundle displacement with the fluid jet unmasked fast adaptation in the current response. (A) At negative potentials, step-like force stimulation (black) resulted in hair bundle creep with no visible fast adaptation in the current. With step-like displacement stimuli (gray), fast adaptation in the current was visible (inset, arrowhead). Similarly, at positive potentials, no fast adaptation was observed with step-like force stimulation (red), but fast adaptation was unmasked with step-like displacements (pink, arrowhead). Displacement was measured in the vertical direction near the middle of the hair bundle at the same location for each stimulus paradigm. Inset traces are expanded to fill the space to better see the onset of the stimuli. (B) Summary of the amount of total adaptation at the 50 ms timepoint between the step-like force and step-like displacement stimulation showed a significant difference between force and displacement stimuli (n=7, −84 mV, *p*=0.000090; n=6, +76 mV, *p*=0.00038). (C) The amount of adaptation at 5 ms would have almost all fast adaptation complete and a small contribution from the slow adaptation process. Adaptation at 5 ms also showed a significant increase indicating the presence of fast adaptation (n=7, −84 mV, *p*=0.000056; n=6, +76 mV, *p*=0.022). (D) The difference in adaptation between the 50 ms and 5 ms time points would be almost completely composed of slow adaptation. This had a significant increase at +76 mV with step-like displacements (n=7, −84 mV, *p*=0.032; n=6, +76 mV, *p*=0.0028). For all adaptation quantifications, step-like displacement responses were not significantly different between negative and positive potentials. (E) Using step-like displacements shortens the time to peak current at both negative and positive potentials (n = 7, −84 mV, *p*=0.0067; n = 6, +76 mV, *p*=0.0038).

With step-like displacements, the amount of adaptation increased significantly. A fast time constant of 1.6 ± 1.2 ms and a slow time constant of 14 ± 3 ms suggests that the total adaptation measured at 5 ms is predominately fast adaptation. We used the difference in total adaptation between the time points of 50 ms and 5 ms to quantify slow adaptation. The amount of total adaptation observed as current decays increased with step-like displacements at 50 ms (Fig. 4B), 5 ms (Fig. 4C), and between 5 and 50 ms (Fig. 4D) for both negative and positive potentials with no significant difference between the two potentials (*p* = 0.61, 0.16, and 0.56, respectively). The time to peak current decreased with displacement stimuli (Fig. 4E), indicating that the current onset is slowed with force stimuli. Across negative and positive potentials, step-like displacements increased the adaptation at 5 ms by 17 ± 8% and the adaptation between 5 and 50 ms by 12 ± 8%, indicating that the step-like displacements led to a greater enhancement of fast current decays.

## Discussion

Our data with the fluid jet are consistent with our previous work using a stiff probe, which demonstrate that fast adaptation is not driven by calcium entry (12). Accounting for hair bundle displacement in three-pulse experiments with fluid-jet stimuli revealed adaptation shifts at positive potentials. Step-like force and displacement stimuli result in different MET current responses. Step-like *force* stimuli results in a mechanical creep with two time constants, which mask the fast current decay associated with fast adaptation. Step-like *displacement* stimuli result in fast current decays at both negative and positive potentials, indicating that calcium entry does not drive fast adaptation.

### Fast adaptation is not driven by calcium entry

After we proposed that calcium entry does not drive fast adaptation (12), others suggested that the results we observed with the stiff probe were an artifact of the stimulus (28). However, we found that PIP_2_ depletion resulted in a loss of fast adaptation with stiff probe stimuli (25), indicating that fast adaptation with the stiff probe data is not a stimulus artifact.

The resting open probability increases with low extracellular calcium or depolarization, and adaptation regulates open probability, which has been taken to argue that lower extracellular calcium and depolarization regulate adaptation. However, mammalian cochlear hair cells can regulate resting open probability by modulating the lipid membrane, independent of fast adaptation (13). Therefore, direct assays are required to test whether calcium entry drives fast adaptation.

We now also demonstrate that data from Corns et al. (2014) using the fluid jet to support calcium-driven fast adaptation can have a different interpretation. Importantly, force and displacement stimuli are not the same. When assaying adaptation with a fluid jet, hair bundle displacement must be quantified because of hair bundle creep. When measuring hair bundle displacement, we observe adaptation that is not driven by calcium entry in three-pulse experiments and that cannot be accounted for by residual slow adaptation (Fig. 3).

We also hypothesize that Corns et al. used an underdamped stimulus that created a time-dependent decrease in current amplitude that appeared to be fast adaptation. We simulated their underdamped stimulus by using an overshoot stimulus (Fig. 2). Under these conditions, fast time-dependent current decays were present at negative but not positive potentials, as with their observation. This result can be mis-interpreted as fast adaptation driven by calcium, when in fact it is due to a stimulus artifact.

Interestingly, an overshoot stimulus can result in current decays at negative, but not positive potentials (Fig. 2B). Initially, we expected the overshoot artifact to cause a similar response at negative and positive potentials. The different responses indicate that there are some properties that are modulated by voltage. Upon further investigation, this result may be due to differences in current rise time. At positive potentials, the current rise time is longer (Fig. 4E). When the overshoot artifact occurs during the longer current rise, it can result in shortening the current rise time and can mask the current decay (see supplemental text), similar to early plateau in the hair bundle displacement using the same stimulus (Fig. 2A onset zoom). Another factor contributing to the lack of fast current decays at positive potentials with an overshoot stimulus is the presence of the slower hair bundle creep component. The slow creep of the hair bundle causes further opening of MET channels, which can reverse any fast current decay and make the current appear to have a stimulus anomaly, rather than being interpreted as a current decay. For example, in Kros et al. (2002), Fig. 2B shows hints of a fast current decay at positive potentials, but the continued current rise due to the slower creep makes the current decay appear like a small notch and not the result of a fast adaptation mechanism.

The most convincing result is the appearance of fast current decays with a fluid-jet stimulus regardless of membrane potential when the driving voltage is modified to achieve a step-like displacement. The step-like displacement most closely mimics the typical step stimuli of a stiff probe. The differences between stimulation methods is complex, such that we do not expect that providing a displacement stimulus with the fluid jet would result in the same exact response we observe with the stiff probe. Notably, the extent of fast adaptation appears greater with the stiff probe (12) than with the fluid jet (Fig. 4A). This discrepancy may be caused by multiple differences in the characteristics of the stimulation methods. Lengthening the stimulation rise time increases the fast adaptation time constant and reduces its extent (10, 27), so differences in stimulus rise times between stimulation methods are likely a major contributing factor. Other contributing factors are the differences in mechanical loading of the hair bundle and pulling of shorter rows of stereocilia due to taller rows moving versus pushing shorter row stereocilia directly. It is unlikely that any one of these is solely responsible for the discrepancy in the adaptation extent. Although the exact responses differ, the basic result remains valid, i.e., that fast adaptation exists at positive potentials and is not driven by calcium entry.

We discuss the Corns et al. (2014) paper because it has the most comprehensive data set supporting calcium-driven fast adaptation; however, other papers have also shown adaptation at negative, but not positive potentials. Indzhykulian et al. used a stiff probe and found that fast adaptation was more prominent at negative than positive potentials (37). These authors assumed the absence of fast adaptation at positive potentials; therefore, they positioned the probe to minimize fast adaptation as a way of avoiding what they assumed was a stimulus artifact (see their results section *Regeneration of MET Occurs in Two Distinct Steps*). Fast adaptation can be saturated even at negative potentials (see Peng et al. (2013), Supplemental Fig. S1). An alternative explanation to their data is that depolarization modulates the saturation point of fast adaptation and that the difference they observed was a voltage-dependent difference between a saturated and unsaturated fast adaptation. Though the correct positioning of a stiff probe is debatable, the fluid jet does not have the same positioning problems, and we still observe fast current decays at positive potentials using a step-like displacement stimuli with the fluid jet (Fig. 4A).

In another paper, Beurg et al. claimed adaptation shifts at negative, but not positive potentials when measuring displacement (31). There are a few concerns with these data and their interpretation. First, the displacement traces do not exhibit the hair bundle creep that we, the authors themselves in another study, and others have observed (Fig. 1C) (28, 33-36, 38). Second, the test pulses after the sustained stimulus do not cover the full range of the activation curve, where not enough negative steps are delivered to determine the minimum MET current after the sustained stimulus. Third, their data show a small adaptation shift at positive potentials. More information about their stimulus and new data with more stimulation steps are needed to resolve this potential discrepancy.

When discussing adaptation, all mechanisms are often considered together, but there are potentially at least 3 mechanisms: fast adaptation, slow adaptation, and lipid modulation. The latter 2 mechanisms are calcium-dependent. Two recent reports claim that in Tmc1 mutant cells (*Beethoven)* with altered calcium permeability, steady-state changes in the activation curve support calcium-dependent adaptation (31, 32). However, neither study directly tests for the calcium dependence of fast adaptation. Corns et al. (2016) performed step stimuli with their fluid jet, which can directly assay fast adaptation (see their Fig. 4-6), but they are likely using an underdamped stimulus. Even so, the data presented have no noticeable effect on fast adaptation. If fast adaptation were driven by calcium entry, then the lowered calcium permeability in the mutant cells would be expected to increase the fast adaptation time constant and/or decrease the extent; however, neither of these occur, which supports that calcium entry does not drive fast adaptation. When interpreting data about the calcium dependence of adaptation, it is important to separate the different mechanisms (fast, slow, and lipid regulation) and to assay each separately on their respective time scales.

### Mechanisms of adaptation

Adaptation is hypothesized to contribute to the hair cell’s dynamic range and frequency selectivity (4, 6); therefore, determining its mechanism is important. Knowing that fast adaptation is not driven by calcium entry is an important step because it narrows the list of possible mechanisms. The manifestation of adaptation differs depending on the mode of stimulation; current decays are observed for displacement steps and hair bundle creep for force steps. This suggests that the hair bundle creep could be a manifestation of fast adaptation (34). One attractive mechanistic model of the creep is viscoelasticity, which is modeled with a spring and dashpot and could indicate a viscoelastic process underlying fast adaptation (14, 39). No direct evidence yet exists in support of a unique mechanism for fast adaptation.

Our data also confirm the presence of a calcium-dependent adaptation mechanism with time constants of ~10 ms. The motor model of slow adaptation proposes that myosin motors dynamically adjust the position of the upper tip-link insertion site to modulate the force transferred to the MET channel depending on the local calcium levels. This model requires a calcium source, namely MET channels, to be localized near the upper insertion site. In mammalian auditory hair cells, the MET channels were localized to the tips of the shorter stereocilia and not near the upper insertion site (40). The channel localization along with the fact that mammalian hair bundles only consist of three rows of stereocilia eliminated the possibility of a direct calcium source modulating myosin motors at the upper tip-link insertion site, therefore questioning the validity of the motor model of slow adaptation (12). Thus, the slow adaptation mechanism may not involve the slipping of myosin motors along the sides of stereocilia. The MET system would still require a tip-link tensioning mechanism in order to have high sensitivity, and myosin motors are the likely candidates for the tensioning mechanism. Myosin motors can still be required to climb up stereocilia to generate tension, but do not regulate the tension directly due to calcium entering through MET channels.

The results in this study confirm that fast adaptation is not driven by calcium entry but that slow adaptation is, reconciling previously conflicting data. The mechanisms of fast and slow adaptation require new molecular models of their function.

## Materials and Methods

### Preparation and recordings

Animals were euthanized by decapitation using methods approved by the University of Colorado IACUC. Organs of Corti were dissected from postnatal day (P) 6-10 Sprague-Dawley rats (a large majority of experiments used P7-P8) of either sex. The tectorial membrane was peeled off, and the tissue was placed in recording chambers as previously described (40). Tissue was viewed using a 60× or 100× (1.0 NA, Olympus) water immersion objective with a Phantom Miro 320s (Vison Research) camera on a SliceScope (Scientifica) illuminated with a TLED+ 525 nm LED (Sutter Instruments). Tissue was dissected and perfused with extracellular solution containing (in mM): 140 NaCl, 2 KCl, 2 CaCl_2_, 2 MgCl_2_, 10 HEPES, 2 creatine monohydrate, 2 Na-pyruvate, 2 ascorbic acid, 6 dextrose, pH=7.4, and 300-310 mOsm. In addition, an apical perfusion, using pipettes with tip sizes of 150-300 µm, provided local perfusion to the hair bundles.

### Electrophysiological recordings

Whole-cell patch-clamp was achieved on the first or second row outer hair cells (OHCs) from the middle to apical cochlear turns using an Axon 200B amplifier (Molecular Devices) with thick-walled borosilicate patch pipettes (2-6 MΩ) filled with an intracellular solution containing (in mM): 125 CsCl, 3.5 MgCl_2_, 5 ATP, 5 creatine phosphate, 10 HEPES, 1 cesium BAPTA, 3 ascorbic acid, pH=7.2, and 280-290 mOsm. For the 0.1 mM and 10 mM BAPTA internal solution, the BAPTA concentration was adjusted accordingly, and CsCl concentrations were adjusted to reach 280-290 mOsm. Experiments were performed at 18-22°C. Whole cell currents were filtered at 10 kHz and sampled at 0.05-1 MHz using USB-6356 (National Instruments) controlled by jClamp (SciSoft Company). Voltages were corrected offline for liquid junction potentials. All experiments used −84 mV holding potential unless otherwise noted. An apical perfusion pipette with extracellular solution was used during patching of hair cells. Only cells with initially < 100 pA of leak were kept for data analysis, except when using 10 mM intracellular BAPTA where < 200 pA was used. Cells also required > 550 pA of initial peak MET current. During a prolonged depolarization, the cell changes resting open probability (12, 13). To have a stable baseline during the positive potential protocols, the cell was depolarized for >10 s to allow the open probability to settle. None of the data presented have had the baseline current zeroed. *Hair bundle stimulation and motion recording*: Hair bundles are stimulated with a custom 3D printed fluid jet driven by a piezo electric disc bender (27 mm 4.6 kHz; Murata Electronics 7BB-27-4L0). Thin wall borosilicate pipettes were pulled to tip diameters of 5-20 micrometers, filled with extracellular solution, and mounted in the fluid-jet stimulator. The piezo disc bender was driven by waveforms generated using jClamp, and the signals were filtered using an 8-pole Bessel filter (L8L 90PF, Frequency Devices Inc.) at 1 kHz and variably attenuated (PA5, Tucker Davis) before being sent to a high voltage/high current amplifier (Crawford amplifier) to drive the piezo disc bender. During stimulations, videos were taken of the hair bundle motion with high speed imaging at 10,000 frames per second using the Phantom Miro 320s at 128 × 128 pixels with a 100 nm effective pixel size. Videos were saved for each stimulation and analyzed offline. Four stimulus presentations were averaged together at each stimulus level unless otherwise stated. In some experiments, a short flexible fiber (long fibers have too much viscous drag) was used to monitor the force of the fluid-jet stimulation. Some fibers were dipped in an oil-based ink to increase fiber contrast. Photodiode measurements were done using a custom-built dual photodiode (SPOT-3D, OSI Optoelectronics) with a custom built onboard differential trans-impedance amplifier (12).

For achieving a step-like hair bundle stimulation with the fluid jet, the voltage step response was fed into a custom-built circuit that mimicked a supercharging circuit with an exponential decay (41). We required two exponential decays to achieve a step-like bundle displacement, so two supercharging circuits were used in series where variable resistors allowed modification of the decay magnitude and time constant to match the observed hair bundle creep. The decay time constants were manually adjusted for each cell to yield a step-like displacement of the hair bundle. These parameters were then fixed for both negative and positive potential protocols for that cell and protocols were taken one after another.

### Hair bundle motion analysis

Custom MATLAB (MathWorks) scripts were used for extraction and analysis of the hair bundle motion. Movie frames were imported into MATLAB and the position of the hair bundle was extracted using a Gaussian fit to a band-pass filtered image (42) for a given vertical row of pixels in the image to yield sub-pixel resolution. For most cells, the vertical displacement of the hair bundle near its center was taken as the displacement of the hair bundle.

### Data analysis

IX plots used the displacement data from the high-speed imaging taking the displacement values at when the peak current occurred for 50 ms step traces. For three-pulse protocols, displacement and current were taken 1 ms after stimulus onset, because the steps were short (5 ms) and peak currents were not always reached during this time period. Normalized currents (I/I_max_) were generated by subtracting the leak current defined as the smallest remaining current during the negative steps and normalizing to the peak current. IX plots were automatically fit using MATLAB with a double Boltzmann equation:

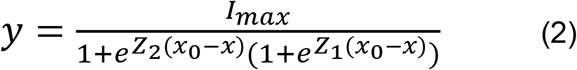

Where Z_1_ and Z_2_ were the slope factors and x_0_ was the set point.

For mechanical stimulus steps, a double exponential decay equation was automatically fit in MATLAB using steps that elicited ~50% maximum current:

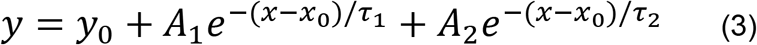

Where τ_1_ and τ_2_ were the decay constants and A_1_ and A_2_ were the respective amplitudes. A_1_/(A_1_+A_2_) is the contribution of fast adaptation and A_2_/(A_1_+A_2_) is the contribution of slow adaptation to the total extent of adaptation. The displacement of the hair bundle was also fit with the same equation starting at the timepoint after the force of the stimulus plateaued, which was 0.5 ms after stimulus onset. Percent adaptation was calculated as (1 – I_steady state_/I_peak_) * 100 for the step that was closest to 50% maximum current.

The time constant calculation based on the adaptation shifts at 10 ms and 50 ms was calculated using the single exponential decay equation. The amount of adaptation occurring at 10 ms is expressed as 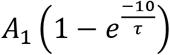 and the amount of adaptation at 50 ms is expressed as 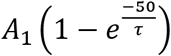 with τ expressed in ms. The ratio of these is equal to 0.475 based on the measurements. This can be rearranged to:

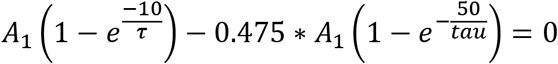

Graphing the left side of the equation and looking for the zero crossings indicates only one zero crossing at τ = 16.7 ms.

All n’s presented were biological replicates of individual cells, where only one cell was used per animal. Each data point was the average of 4 presentations of the same stimulus intensity (technical replicates), unless otherwise noted.

Data were analyzed using jClamp, MATLAB (MathWorks), and Excel (Microsoft). Graphs were created using MATLAB and Adobe Illustrator. The mechanosensitive current/maximum mechanosensitive current was used as P_open_, where we assume an observable P_open_ of 100%. The maximum MET current was the difference between the current values elicited from the maximal negative and maximal positive stimulation.

Statistical analysis used Student’s two-tailed t-tests from MATLAB or Excel (Microsoft) unless otherwise stated. Paired tests were used when comparing across data points in the same cell, and unpaired-unequal variance tests were used to compare data across cell populations. Significance (*p*-values) were as follows: * *p*<0.05, ** *p*<0.01, *** *p*<0.001, **** *p*<0.0001. Data were presented as mean ± standard deviation unless otherwise noted.

## Acknowledgements

Work was supported by R00 DC013299 and R01 DC016868 to AWP, by R01 DC003896 to AJR, and by core grant P30-44992. AM was supported by T32 NS099042. We thank Kurt Beam, Artur Indzhykulian, and David Corey for critical feedback of this manuscript.

