## Supplemental Information for "Calcium entry does not drive fast mechanotransduction adaptation in cochlear hair cells"

*Supplemental Text*

With the hair-bundle creep masking the fast current decays, the current rise time could lengthen. Therefore, a cell with a long current rise time when using a step-like force stimulus would have a shorter current rise time with a step-like displacement stimulus (Supplemental Fig. S2A left). The times to peak current were significantly shorter for step-like displacements of the hair bundle when compared to step-like force stimuli, which further supports that the hair-bundle creep masks the fast current decays (Fig. 4E). For step-like displacements, there was also an inverse correlation between the total adaptation at 5 ms and the time to peak current (Supplemental Fig. S2B). This indicates that a cell with a short current rise time was more likely to exhibit current decays (Supplemental Fig. S2A). The inverse correlation may account for the non-significant differences between the total adaptation at 5 ms between negative and positive potentials for step-like displacements (Fig. 4C), since at negative potentials, shorter times to peak current were observed (Fig. 4E).

*Supplemental Figures*

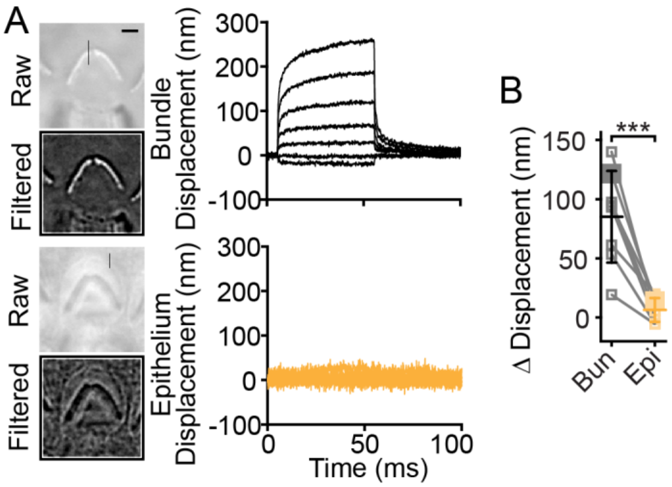


**Fig. S1.** Hair-bundle creep was not due to epithelium movement. (A) Stimulation of the hair bundle led to hair-bundle creep (black traces) that was not visible in the epithelium movement (orange traces). Raw and filtered images are shown with the location of measured motion shown with the vertical black line in the raw image. Horizontal scale bar = 2 µm. (B) Summary of the change in displacement of the hair bundle (Bun) or epithelium (Epi) from 0.5 ms after stimulus onset to the end of the stimulus step for the largest stimulation is plotted (n=8, $p$=0.00045). Lines connect data from the same cell with the same stimulus intensity. Data from panel A are shown with larger symbols.

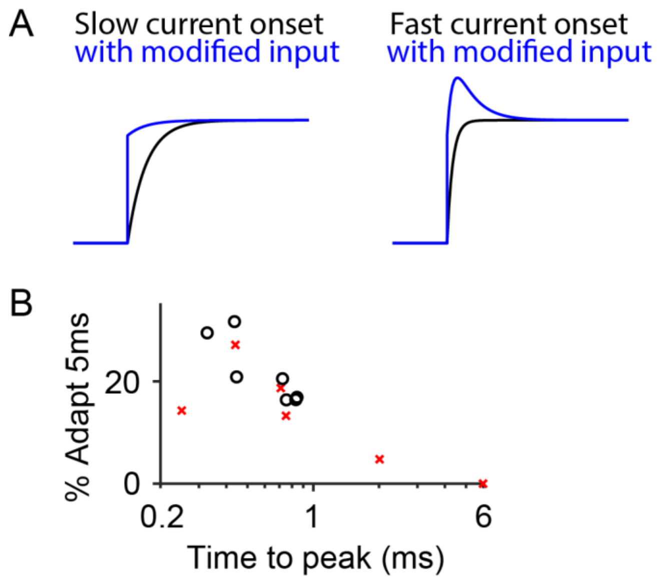


Supplemental Fig. S2. Effect of current rise time on the measured adaptation percentage. (A) With a long current rise time, adding an exponential decay function to the curve may only shorten the current rise time (left), but adding the same exponential decay function to a short current rise time results in current decay. The exponential decay function is similar to what the modified stimulus may do to achieve a step-like displacement. (B) Plot of the percent adaptation at 5 ms vs. the base 10 log of the time to peak current for step-like displacement stimulation at negative (black circle) and positive potentials (red x) indicates a negative correlation between these parameters. Pearson's correlation coefficient = -0.80, $p = 0.0011$.
